# An Advancing Front of Old Age Human Survival

**DOI:** 10.1101/361782

**Authors:** Wenyun Zuo, Sha Jiang, Zhen Guo, Marcus Feldman, Shripad Tuljapurkar

## Abstract

Old age mortality decline has driven recent increases in lifespans, but there is no agreement about trends in the age-pattern of old deaths. Some hypotheses argue that old-age deaths should have become compressed at high ages, others that old-age deaths should have become more dispersed with age, and yet others are consistent with little change in dispersion. However, direct analyses of old-age deaths presents unusual challenges: death rates at the oldest ages are always noisy; published life tables must assume an asymptotic age pattern of deaths; and the definition of “old age” changes as lives lengthen. Here we use robust percentile-based methods to overcome these challenges and show, for 5 decades in 20 developed countries, that old-age survival follows an advancing front, like a traveling wave. The front lies between the 25th and 90th percentiles of old-age deaths, advancing with nearly constant long-term shape but annual fluctuations in speed. The existence of this front leads to several predictions that we verify, e.g., that advances in life expectancy at age 65 are highly correlated with the advance of the 25th percentile, but not with distances between higher percentiles. Our unexpected result has implications for biological hypotheses about human aging, and for future mortality change.

---

Longer lives are an achievement (*1*) and a challenge (*2*,*3*). Recent increases in human longevity are driven by reductions in post-retirement (ages 65+) mortality (*4*) but the age-pattern of old deaths is hotly debated. If lifespans are approaching a limit (based on biological hypotheses (*5*), or the oldest ages at death (*6*)), death should have become compressed towards the highest ages. If long life depends on endowments (economic (*7*), or genetic (*8*)), the oldest deaths should over time become distanced from earlier deaths. But if long life is driven by medical advances (*9*) and decreasing old-age disability (*10*), there may be little or no change in the age-pattern of old deaths. However, direct analyses of old-age deaths face unusual challenges. First, in any year the oldest ages are always reached by the fewest survivors so corresponding death rates are noisy. Second, published life tables assume a maximum age and age-pattern of oldest deaths, even though neither is known (*11*). Third, the definition of “old age” is changing: e.g., Japanese females had an 80% probability of living past age 60 in 1960, past age 70 by 1977, and past age 80 by 2011 (*11*). Partly as a result, previous work has focused on the compression of adult deaths (*12*), deaths near a modal adult age (*13*,*14*), or on oldest death records (*15*). Here we use percentiles (extending (*12*)), to overcome these problems and robustly examine the age-pattern of old-age mortality.

Annual life tables (female, male, 1960 to 2010 (*11*)) provide age-specific death rates. To focus on old-age deaths we consider individuals in each year who are alive at age 65 and thereafter experience death rates for that year: the age by which *q*% of such individuals would die is called *A_q_* (*16*). The ages *A_q_* are percentiles of the period death distribution (with *A*_0_ = 65), and describe the shape of old-age deaths (Fig. 1A). E.g., percentiles for a narrow distribution are closer together than for a wider distribution. Given *q*, the corresponding *A_q_* is computed forward from age 65 and so is unaffected by later deaths. To minimize noise we focus on percentiles up to *A*_90_, a high age (e.g., for the US in year 2000, *A*_90_ is 94.5 for females, and 91.4 for males).

**Figure 1:**
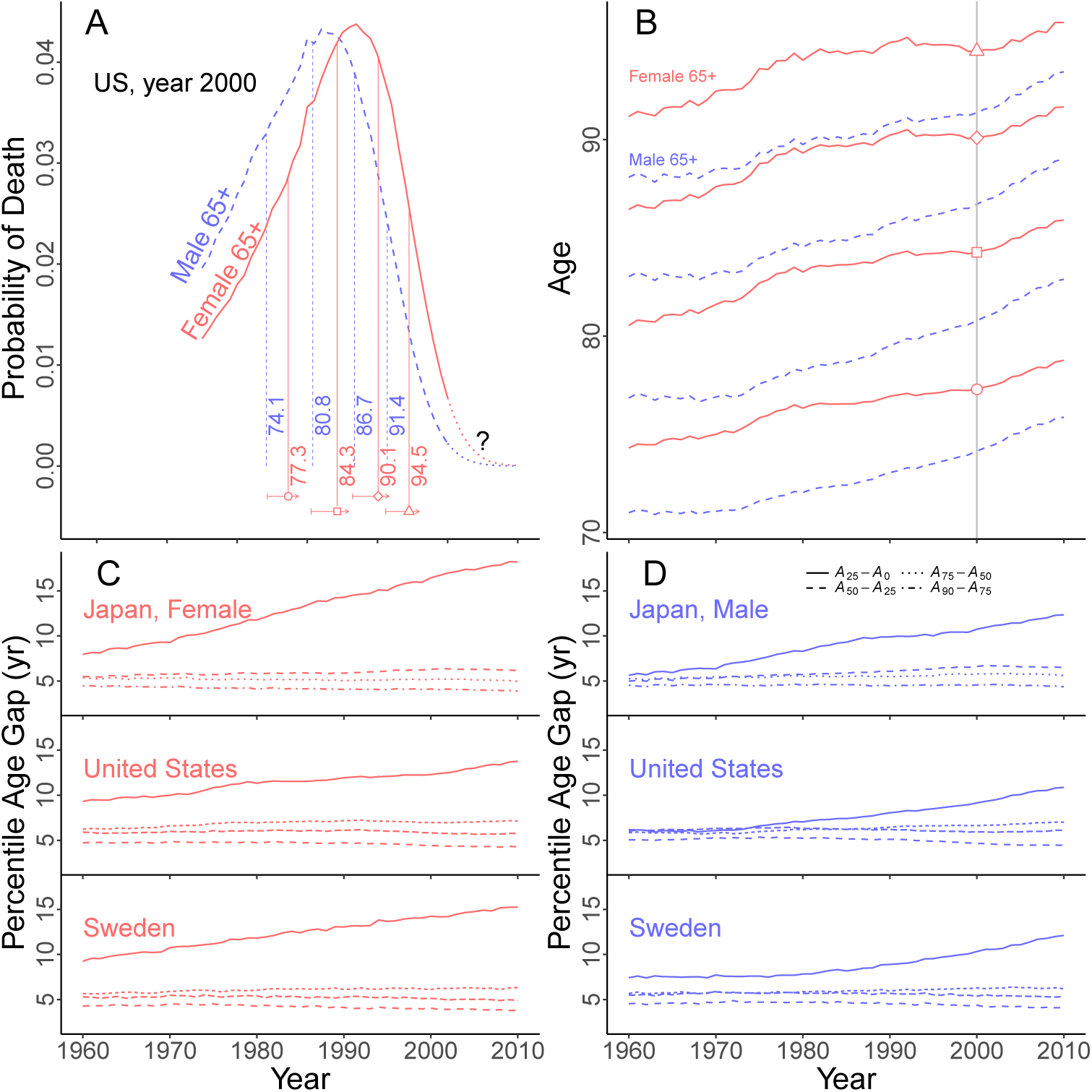
**A.** Probabilities of death at ages past 65 for US females, red solid, and males, blue dashed, using death rates in year 2000. Question mark indicates oldest ages with uncertain death rates. Vertical (solid, dashed) lines mark corresponding 25th, 50th, 75th and 90th percentiles of death. Horizontal solid arrows show movements from 1960-2010 of the *A_q_*; symbols on arrows mark year 2000. B. Changes in *A*_25_, *A*_50_, *A*_75_, *A*_90_ for US females, red solid, and males, blue dashed; symbols mark year 2000. C and D. Interval (*A*_25_ – 65), and between consecutive percentiles *A_q_* (for q =25, 50, 75, 90) for Japan, the US and Sweden (top to bottom panels). C, females (red); D, males (blue). Only (*A*_25_ – 65) rises steadily. All other intervals show little long-term trend and small annual fluctuations.

All age percentiles increased for US females over 5 decades (Fig. 1B), albeit with annual fluctuations. But surprisingly, the age intervals between adjacent percentiles appears nearly constant, implying that the period distribution of old-age female deaths had nearly the same shape for 5 decades. Compared to females, males die earlier (Fig. 1A) with age percentiles changing at different rates (Fig. 1B), but here too age intervals between adjacent percentiles appear nearly constant for 5 decades. These observations led to a comparison of Japan, Sweden and the US (Fig. 1, C and D). In each country and sex, the age percentile *A*_25_ moves steadily away from age 65, but intervals between adjacent higher percentiles change little and have only small annual fluctuations, suggesting a stable shape for old-age deaths.

Next, we examined annual speeds (change over a year), and long-term speeds (the average of the annual speeds from 1960 to 2010) for age percentiles *A*_1_, *A*_2_,…, *A*_99_ (Fig. 2, A and B; figs. S1 to S20 (*16*)). Percentiles between A_25_ and A_90_ have similar positive long-term speeds. But annual speeds are highly variable, especially at older ages (Fig. 2, A and B), which should imply high annual variability of the intervals between consecutive percentiles. However, the latter intervals display only small annual fluctuations (Fig. 1, A and B). Therefore, even when adjacent percentiles each move by a large amount, the distance between them stays nearly constant – implying a survival front with nearly constant shape. So we must find a strong positive correlation between annual changes in consecutive percentiles, which we find in Japan [(Fig. 2, C and D) and elsewhere (figs. S21 to S39). The long-term speeds of intervals between consecutive percentiles can be positive or negative but are small, e.g., for negative speeds, (*A*_90_ – *A*_75_) would take decades to decline by 20% (table S1).

**Figure 2.**
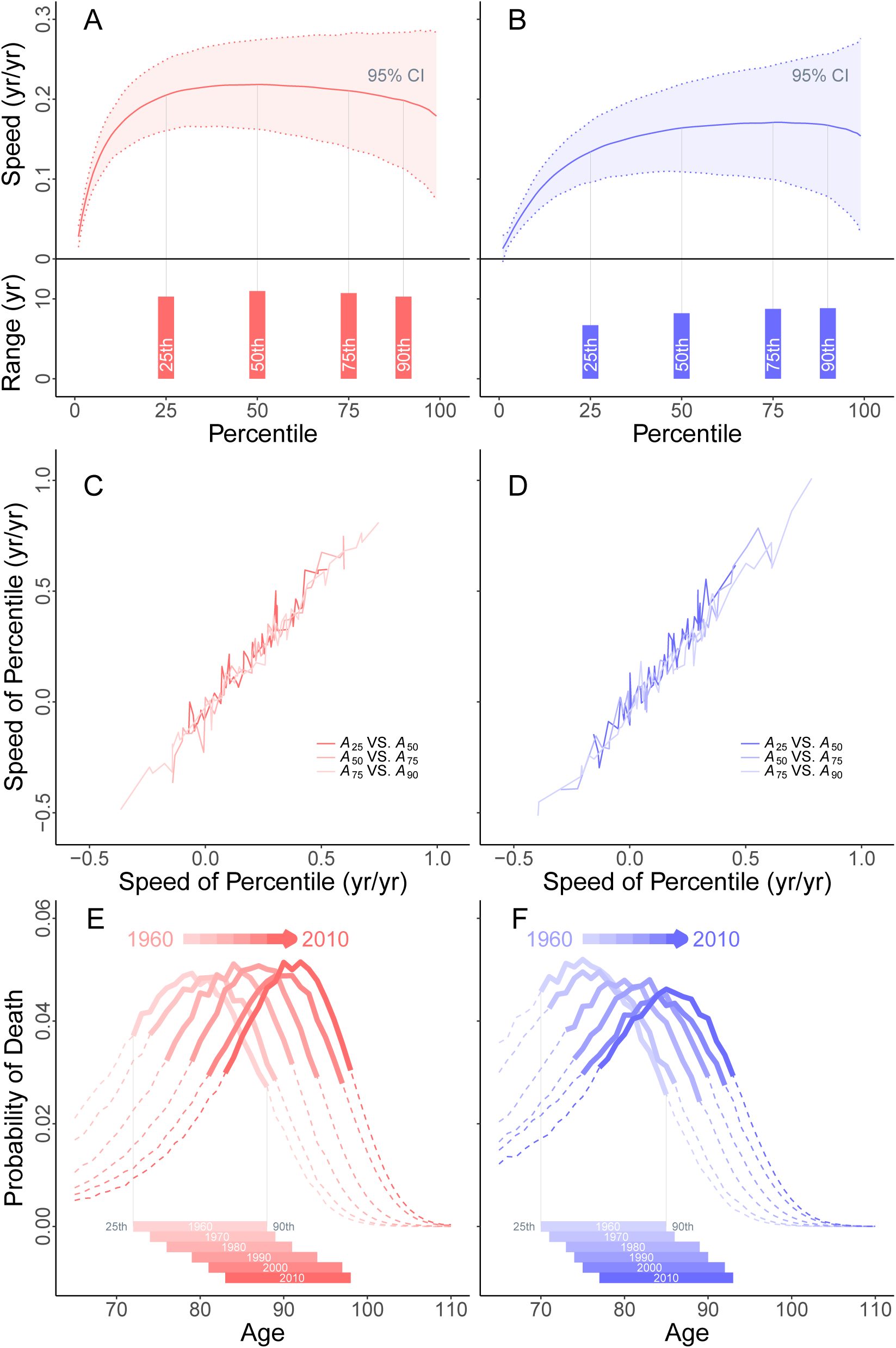
*(preceding page)*: An example from Japan.**A** and **B**. Speeds (rates of movement) for percentiles *A_q_* at every 1% (using raw data on death ages above *A*_90_). Top, speeds for percentiles for females on left, respectively males on right. Solid line (red, respectively blue) indicates long-term speed. Percentiles from the 25th to the 90th show similar long-term speeds (note the vertical scale). Long-term speeds 0.2 yr/yr for females. Also shown: the 95% confidence interval (1.96 x Std Dev, distributions symmetric and approximately normal) for annual speeds: dotted lines and bands (pink, respectively blue). Annual variability is high (compare modest annual variability for intervals between percentiles, Fig. 2A). Bottom panels, solid bars (red, females, blue, males) show ranges. **C** and **D**. For each ending year, annual speeds of percentiles; left, females in red; right, males in blue. Annual change in *A*_50_ on the vertical versus annual change in *A*_25_ on the horizontal, and correspondingly for the pairs *A*_75_, *A*_50_ and *A*_90_, *A*_75_. **E** and **F**. Probability distributions of deaths at 65+. Solid line, advancing front of old-age survival (between **A**_25_ and **A**_90_, dashes outside that range) for decades 1960-2010. Left, females in red; right, males in blue. For each year, solid bars, bottom, show distance between **A**_25_ and **A**_90_.

These results strongly support an advancing old-age survival front with a nearly stable shape between the 25th and (at least) the 90th death percentiles (Fig. 2, E and F for Japan). The shape fluctuates modestly over time but with no long-term trend. The male front has lower long-term speed and is more dispersed than the female front. We found old-age survival fronts for females and males in 20 industrialized countries (*11*) over the 5 decades.

The existence of an old-age survival front with nearly constant shape yields four testable predictions. First, annual changes in the *A_q_* should be strongly positively correlated across successive percentiles – as verified earlier (Fig. 2, C and D).

Second, the life expectancy *e*_65_ at age 65 should increase as the front advances. Thus, across countries, the rate of increase of *e*_65_ should be positively correlated with the long-term speed of (*A*_25_ – 65), but uncorrelated with changes in later intervals, such as (*A*_75_ – *A*_50_). These predictions hold for for both sexes (Fig. 3 and figs. S40 and S41).

**Figure 3:**
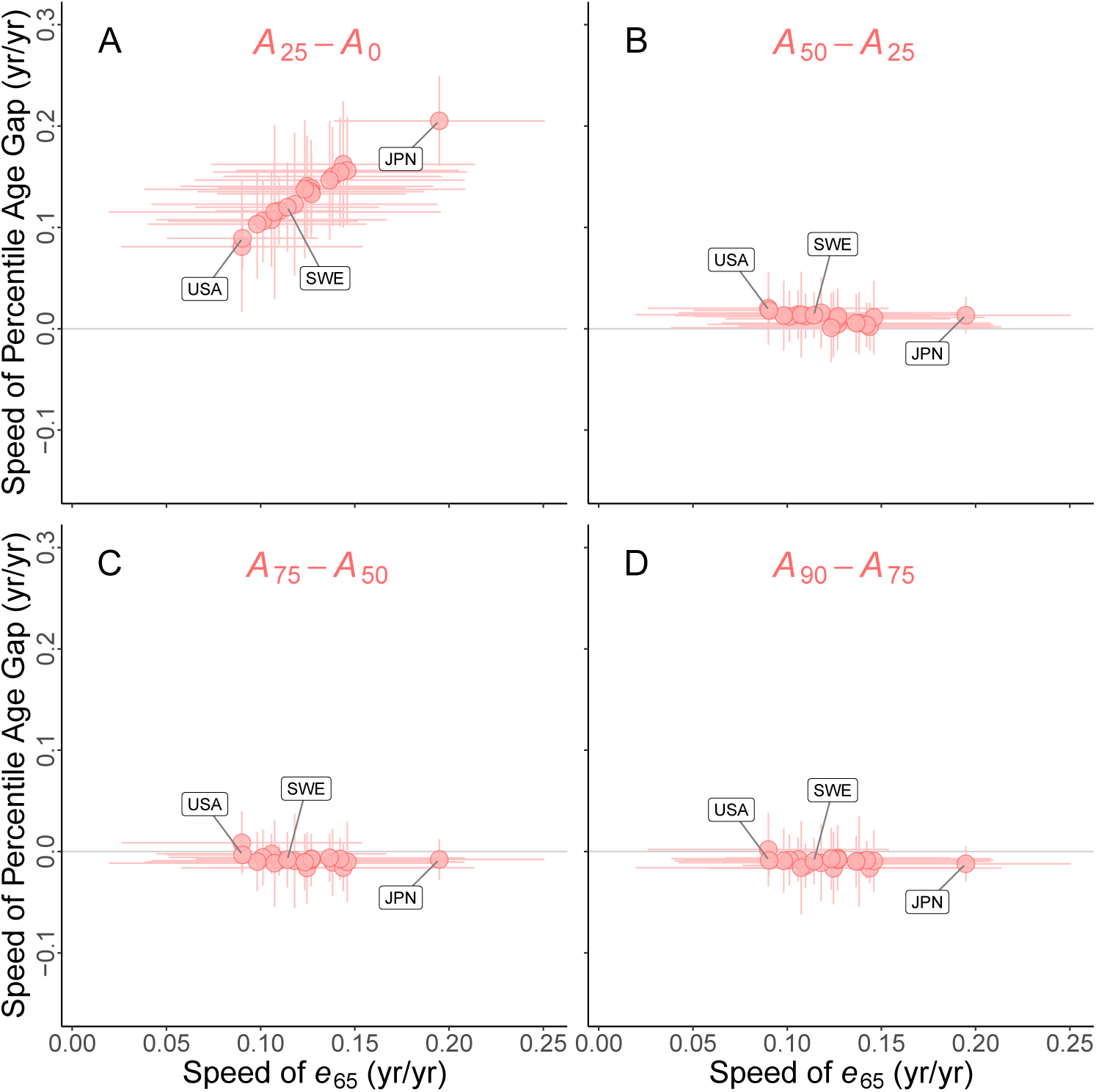
**A**. Vertical: long-term speed of (*A*_25_ – 65); horizontal: long-term speed of *e*_65_ (life expectancy at age 65). Each dot is a country; for each dot, the vertical and horizontal lines show 95% confidence intervals for annual speeds. There is a strong positive correlation. **B**. Vertical: long-term speed of (*A*_50_ – *A*_25_); horizontal: long-term speed of *e*_65_; each dot is a country. There is nearly zero correlation. **C**. Long-term speed of (*A*_75_ – *A*_50_) (vertical) uncorrelated with longterm speed of *e*_65_ (horizontal axis). **D**. Long-term speed of (*A*_90_ – *A*_75_) (vertical) uncorrelated with long-term speed of *e*_65_ (horizontal).

Third, the variability of deaths after age 65 should increase over time. Here, for any age *a*, variability among later deaths is measured by the standard deviation *s_a_* of ages of death (*13*). This prediction, implied by the steady movement of the survival front away from age 65, holds (Fig. 4, A and B; figs. S42 and S43), as was shown, though not explained, in (*17*).

**Figure 4:**
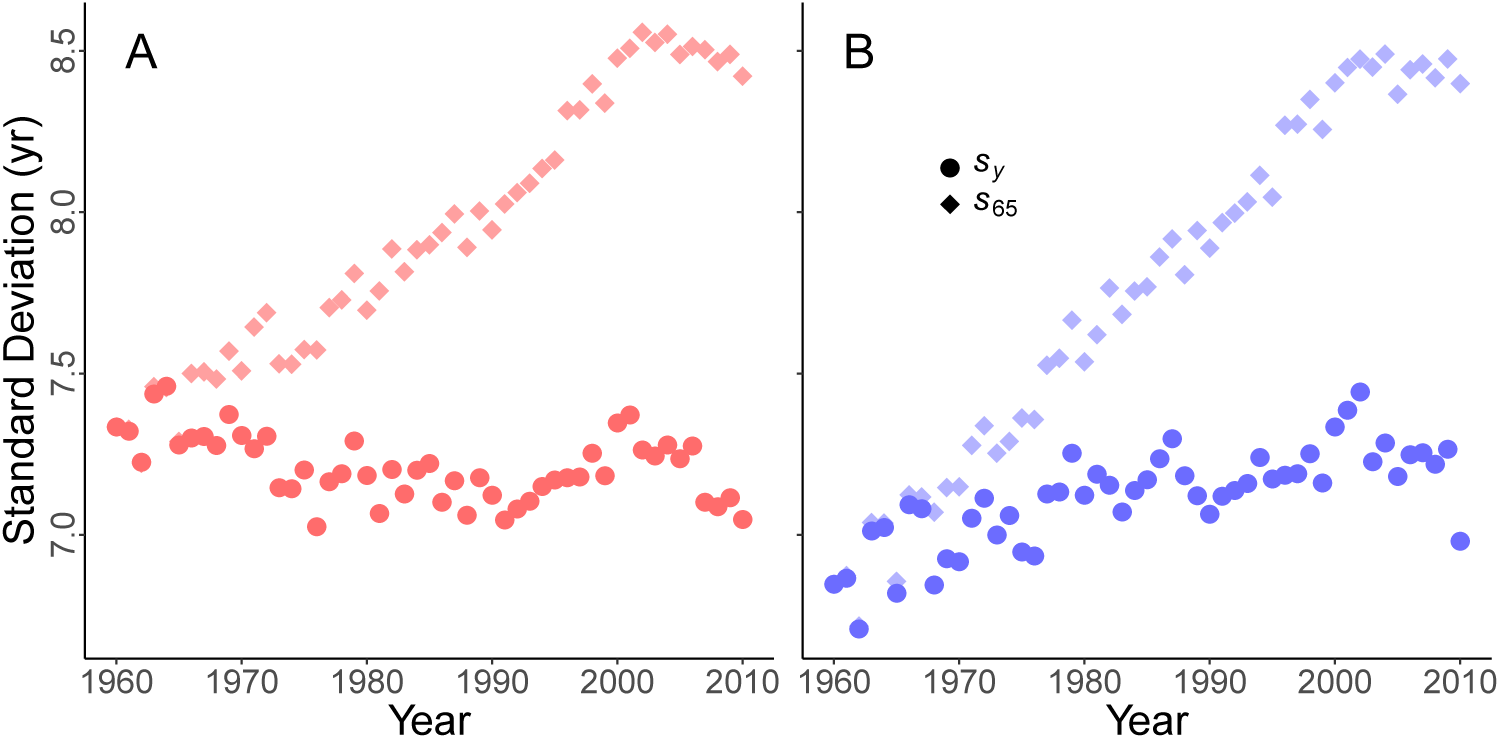
**A** and **B**. Vertical: variability (measured by standard deviation of age at death) for deaths past age 65 (diamonds) or a moving age *y* (circles). Horizontal: year from 1960 to 2010. Left, Japanese females, red; right, Japanese males, blue. Diamonds: rapid increase of *s*_65_. For each year *t*, moving age *y* = 65 + *v*(*t* – 1960), with *v* the long-term speed of the survival front. Circles show variability *s_y_*.

Fourth, there should be no increase, or even a decline, in the variability of deaths that occur past an age that moves along with the survival front. For each year *t* from 1960 to 2010, define age *y* = 65 + *v* (*t* – 1960), where *v* is the long-term speed of the survival front (measured here by the long-term speed of *e*_65_). We predict that deaths after age *y* have a near-constant, or even declining, dispersion *s_y_*, in sharp contrast to the increasing trend in *s*_65_. We find such a contrast for Japan (Fig. 4, A and B), and for all countries (figs. S42 and S43).

We conclude that an advancing old-age front characterizes old-age human survival in 20 developed countries. Our findings provide no support for a limit to human lifespan, certainly not at an age that affects the survival front now or for many decades. Nor do our results suggest that endowments, biological or other, are a principal determinant of old-age survival. Our result is consistent with increases in the age of transition to disability, and with the hypothesis that deaths result from an accumulation of detrimental changes at a rate influenced by prosperity.

Note that the location of the survival front in any year may be affected by earlier death, e.g., due to opioids (*18*), as reflected in the annual volatility of changes in the percentiles (Fig. 2, C and D). The advance in survival that we find suggests that the effects of inequality on mortality (*19*) may be much smaller at old ages than among younger adults. Our results can be used to bound the parameters of some mortality models, but do not explain differences in the location and speed of the front between sexes or countries. The surprising regularity we report should be used to improve mortality forecasts (*20*), and implies that we must welcome continued aging in spite of its challenges.

## Acknowledgements

We thank the Morrison Institute for Population and Resource Studies at Stanford University and the School of Sociology, Huazhong University of Science and Technology, for support of S.J. Useful comments were provided by Peter Cameron, Pradip Rathod, Diana Rypkema and Shubha Tuljapurkar.

## Author contributions statement

S.T. conceived the analyses, S.J. and W.Z conducted the analyses, S.T. and W. Z. wrote the paper with substantial help from S.J., and input from M.F. All authors reviewed the manuscript.

## References

1. J. W. Vaupel, Nature 464, 536 (2010).

2. R. Lee, J. Skinner, The Journal of Economic Perspectives 13, 117 (1999).

3. J. Bongaarts, Population and Development Review 30, 1 (2004).

4. R. Lee, S. Tuljapurkar, Demography 34, 67 (1997).

5. T. B. Kirkwood, S. N. Austad, Nature 408, 233 (2000).

6. X. Dong, B. Milholland, J. Vijg, Nature 538, 257 (2016).

7. J. Poterba, S. Venti, D. A. Wise, Working Paper 21682, National Bureau of Economic Research (2015). http://www.nber.org/papers/w21682.

8. J. Kaplanis, et al., Science 360, 171 (2018).

9. D. Goldman, et al., American journal of public health 99, 2096 (2009).

10. E. Crimmins, H. Beltrán-Sánchez, The Journals of Gerontology Series B: Psychological Sciences and Social Sciences 66, 75 (2011).

11. University of California, Berkeley (USA), Max Planck Institute for Demographic Research (Germany), Human mortality database, http://www.mortality.org or http://www.humanmortality.de. Accessed: 2017-08-30.

12. J. R. Wilmoth, S. Horiuchi, Demography 36, 475 (1999).

13. R. Edwards, S. Tuljapurkar, Population and Development Review 31, 645 (2005).

14. A. R. Thatcher, S. L. K. Cheung, S. Horiuchi, J.-M. Robine, Demographic Research 22, 505 (2010).

15. V. Kannisto, Demographic Research 3 (2000).

16. See the supplementary materials.

17. M. Engelman, V. Canudas-Romo, E. M. Agree, Population and Development Review 36, 511 (2010).

18. A. Case, A. Deaton, Proceedings of the National Academy of Sciences 112, 15078 (2015).

19. R. Chetty, et al., Jama 315, 1750 (2016).

20. S. Tuljapurkar, N. Li, C. Boe, Nature 405, 789 (2000).

